# Hematophagy generates a convergent genomic signature in mosquitoes and sandflies

**DOI:** 10.1101/2024.08.07.607008

**Authors:** Julien Devilliers, Ben Warren, Ezio Rosato, Charalambos P. Kyriacou, Roberto Feuda

## Abstract

Blood-feeding (hematophagy) is widespread across Diptera (true flies), yet the underlying molecular mechanisms remain poorly understood. Using phylogenomics, we show that four gene families associated with neuro-modulation, immune responses, embryonic development, and iron metabolism have undergone independent expansions within mosquitoes and sandflies. Our findings illuminate the underlying genetic basis for blood-feeding adaptations in these important disease vectors.

## Introduction

Insect vectors play a pivotal role in the transmission of major human diseases, including dengue, zika, malaria, and leishmaniasis, resulting in over 700,000 deaths annually (WHO, 2020). Hematophagy (blood-feeding) is largely responsible for the transmission of diseases and is widely distributed among Insecta, likely evolving independently multiple times (Mans, 2011). This behaviour requires a suite of adaptations, including specialised biting structures to access blood, the release of anti-coagulants and host-immune modulators plus the ability to withstand high levels of toxic metallic ions present in blood (Barillas-Mury *et al*., 2022). Recent genomic analyses carried out on a few hematophagous insects have suggested that the expansion of gene families implicated in heat-shock responses and chemosensory function may constitute a distinctive genetic signature associated with blood-feeding (Freitas and Nery, 2020). However, a comprehensive understanding of the genomic underpinnings of hematophagy remains constrained, partly due to the paucity of genomic resources.

The Diptera contribute many blood-feeding disease vectors, including mosquitoes, sandflies, blackflies and tsetse flies. Moreover, the wealth of high-quality genomic data available for this order provides an opportunity to investigate genomic changes associated with hematophagy, particularly in mosquito and sandfly species. Here, we identify five gene families that appear to have evolved at different rates in mosquitoes and sandflies compared to other dipteran species. We suggest that these modifications may provide an indicator of the genetic changes occurring during the evolution of hematophagy.

## Results

### Phylogenetic inference and ancestral state reconstruction support the independent evolution of blood-feeding

The initial crucial step to investigate genomic changes linked to blood-feeding is the construction of robust phylogenetic trees. We developed a comprehensive dataset of proteomes from 64 dipteran species, maximising taxonomical diversity and proteome completeness, and including representatives of mosquitoes (Culicidae) and sandflies (Phlebotominae) (see Table S1). We then inferred phylogenetic relationships combining two approaches (see Methods): a **(1)** Maximum Likelihood (ML) inference on a single-copy gene supermatrix and a **(2)** gene trees-based method taking into account for incomplete lineage sorting (*i*.*e*., remaining gene flow among species after speciation).

The trees obtained using these two approaches are largely consistent, supporting the monophyly of Culidicae (UltraFast bootstrap, UFB=100 and Local Posteriori Probability=1.00) and Phlebotominae (UFB=100 and Local Posteriori Probability=1.00). However, the two methods identified some incongruences (Figure S1) restricted to the phylogenetic position of *Simulium sp*. (Simuliidae), *Dixa sp*. (Dixidae) and *Corethrella calathicola* (Corethrellidae).

Based on taxonomical distribution, it has been suggested that hematophagy emerged independently multiple times in Diptera (Mans, 2011; Wiegmann *et al*., 2011). To test this more formally, we used an ancestral state estimation method using a phylogenetic ridge regression (see Methods) on both trees (Figure 1 & S2). Our analyses (Figure 1 & S2) suggest that the last common ancestor of our tree was non-blood-feeding, while the last common mosquito ancestor (LCMA, node A in Figure 1) and last common sandfly ancestor (LCSA, node B in Figure 1) were blood-feeding with probabilities of 0.67 and 0.97, respectively. Such results support the hypothesis of behavioural convergence between the two groups.

**Figure 1.**
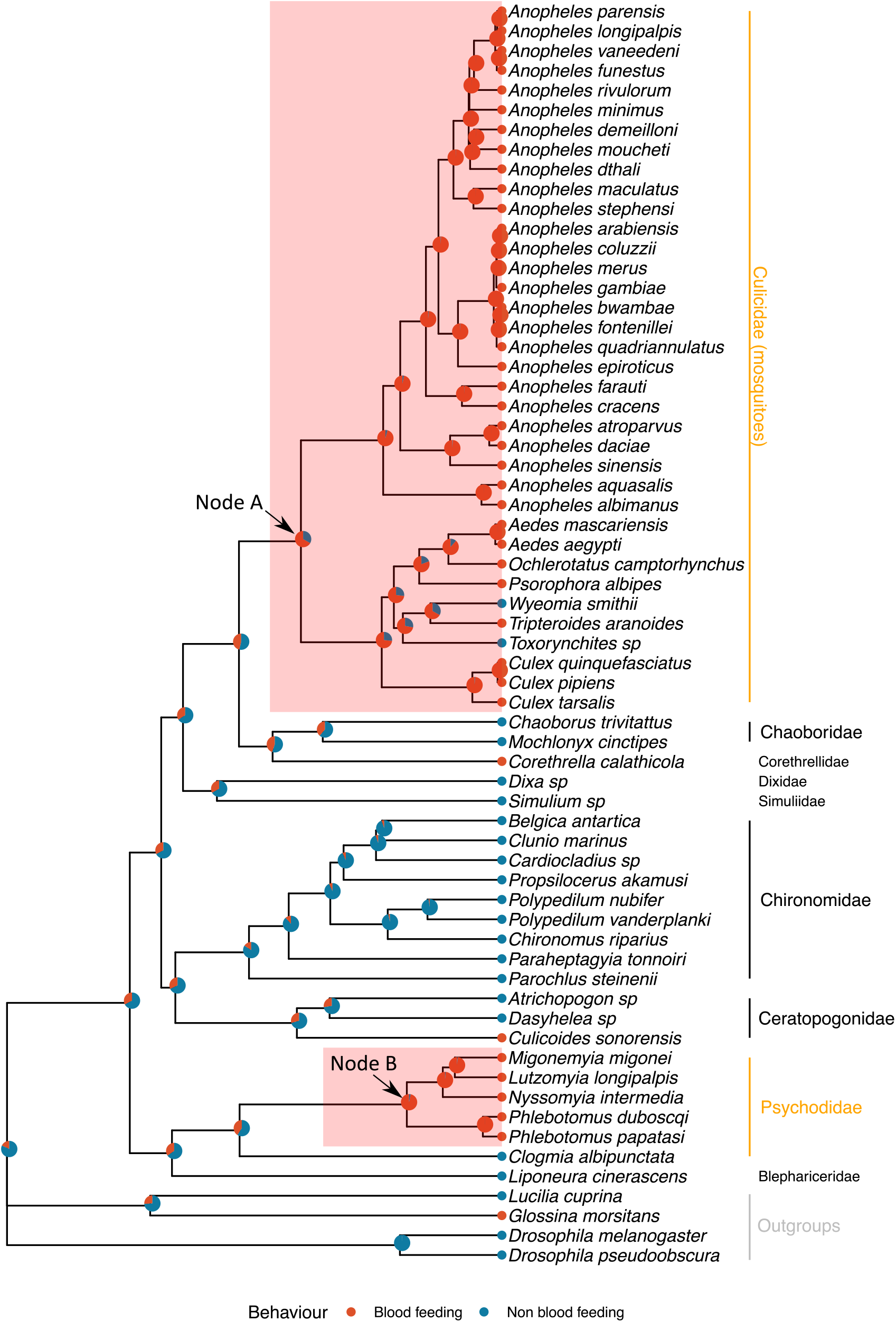
Phylogenetic and ancestral trait inference of blood-feeding behaviour in Culicomorpha and Psychidomorpha. The ultrametric ASTRAL tree was obtained from 63,621 gene trees with ASTRAL-Pro (Zhang et al., 2020) and divergence times computed in r8s (Sanderson, 2003). The colour of the nodes shows the probabilities for ancestral behaviour (blood feeding [red] or non-blood feeding [blue]) from a phylogenetic ridge regression obtained with RRphylo (Castiglione et al., 2020). Nodes A and B, respectively, represent our node of interest with the emergence of hematophagy in mosquitoes and sandflies.

### Genomic signature of convergence in hematophagy

A leading hypothesis proposes that the emergence of novel traits correlates with the expansion and contraction of gene families (*i*.*e*., orthogroups [OGs]) during evolution (Kaessmann, 2010; Osipova *et al*., 2023). We modelled OGs dynamics using a birth-death model to infer evolutionary dynamics related to gene family evolution (see Methods). Such methodology allows for the identification of significant changes in the rate of evolution of OGs against the average rate of evolution per genome across all species in the tree.

Independently from tree topologies, we observed a change in the rate of evolution of 70 OGs in sandflies, with 61 OGs showing an expansion and 9 a contraction (p<0.05). In contrast, in mosquitoes, we observed an expansion of 261 OGs (p<0.05). Among the above OGs, we have identified five OGs represented in both clades. The *neuronal calcium-binding protein* OG (OG0000128) showed an expansion within mosquitoes and a contraction within sandflies. The four other OGs, *octopamine receptors* (OG0001113), *TBP-associated factor 3* (OG0001405), *qin* (OG0001557) and *WD-repeat protein outer segment 4* (OG0002006) exhibit an increased diversification in both clades (Figure 2 & Supplementary Materials 1.0 S1).

**Figure 2.**
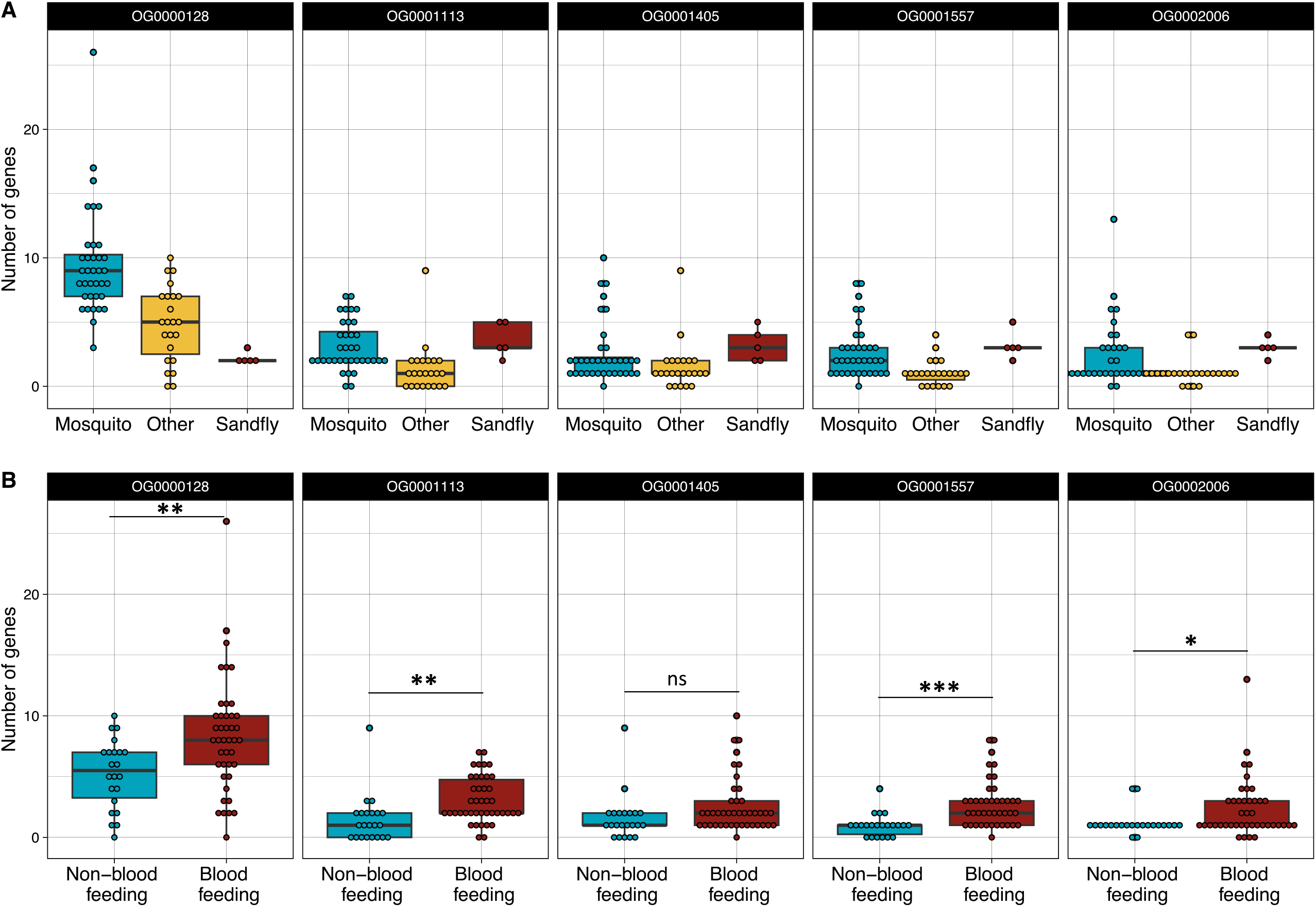
Boxplot showing the repartition of number of genes in gene families showing a modification of their rate of evolution in mosquitoes and sand-flies. A - Distribution of number of genes in mosquitoes, sand-flies and other species from our dataset. B - Distribution of number of genes in blood feeding and non-blood feeding species. An ANOVA have been performed to test the effect of hematophagy on gene copy number for each OG: *, p<0.05; **, p<0.01; ***, p<0.001; ns, p>0.05.

To further clarify the evolutionary dynamics of these OGs, we performed a gene-tree-to-species-tree reconciliation, taking into account the protein sequences (see Methods). These analyses suggested that within the *neuronal calcium-binding protein* family (OG0000128) three gene duplications occurred in the LCMA and three gene losses in the LCSA. We identified one and two duplications, respectively in the LCMA and LCSA, for *octopamine receptors* (OG0001113), *TBP-associated factor 3* (OG0001405), *qin* (OG0001557) and *WD-repeat protein outer segment 4* (OG0002006) gene families (Figure S3 & S4). When comparing blood-feeding with non blood-feeding species in our dataset, we observed that hematophagy correlates with a higher number of paralogs than non-biting species in all OGs (OG0000128: F=7.828, p=0.0068; OG0001113: F=10.344, p=0.0021; OG0001557: F=16.492, p=0.0001; OG0002006: F=4.6255, p=0.0354) except *TBP-associated factor 3* (OG0001405: F=3.2269, p=0.0773) (Figure 2B).

## Discussion

Hematophagy is widespread among arthropods and has emerged independently multiple times (Mans, 2011). Even if differences exist in the physiological pathways used for biting, the general mechanism remains similar among groups of arthropods (Barillas-Mury *et al*., 2022). The transition to blood feeding is challenging as multiple barriers must be overcome; for instance, host localisation, access to their blood, digestion and detoxification of the blood meal being the most evident. A simple signature of gene family evolution is represented by variation in the number of gene copies over time: duplication of a gene can lead to different selective regimes on both copies of the gene, allowing for the evolution of novelties (Kaessmann, 2010). Similarly, gene loss has been shown to allow for adaptation by improving metabolic efficiency (Osipova *et al*., 2023).

In hematophagous mosquitoes and sandflies, we have identified five gene families showing evolutionary changes in the number of paralogs present in the last common ancestor of each group. The two clades occupy different ecological niches; for example, mosquito larvae are aquatic while sandfly larvae grow in soil (Cecílio *et al*., 2022). Furthermore, while adults from both groups feed on sugars, sandflies are more flexible, foraging on fruits in addition to nectar from flowers (Junnila *et al*., *2011*). Their only commonality is the hematophagy of adult females for oviposition. We might thus expect a genetic signature of hematophagy with either diversification or contraction of the same gene families in both clades.

We identified 5 OGs with significant changes in their gene copy dynamic that is associated with the emergence of hematophagy in mosquitoes and sandflies: *neuronal calcium-binding protein* (OG0000128), *octopamine receptors* (OG0001113), *TBP-associated factor 3* (OG0001405), *qin* (OG0001557) and *WD-repeat protein outer segment 4* (OG0002006). Our results differ from previous studies that used a few representatives of some biting species among insects, which detected a contraction of the genes encoding chemosensory proteins and an expansion of those encoding carboxylic ester hydrolases (Freitas and Nery, 2020). However, the methodology of both studies differs, with a focus in our work on closely related species from two groups of hematophagous dipterans, against a few representatives from orders among Insecta in Freitas and Nery (Freitas and Nery, 2020). Furthermore, different from Freitas and Nery (Freitas and Nery, 2020) statistical thresholds in our studies are higher, targeting expansion and contraction events common to the full clade and not simply in two hematophagous species from different biting lineages.

The *neuronal calcium-binding protein* gene family (OG0000128) shows increased diversification in mosquitoes but contraction in sandflies (Figure 2A). In mosquitoes, this gene family includes hippocalcin, a brain-specific protein involved in sleep (Chen *et al*., 2019); frequenin, involved in synaptic exocytosis, neurite outgrowth and neuronal specification during embryonic development (Dason *et al*., 2012; Pongs *et al*., 1993); calcenilin, a voltage-gated potassium channel involved in calcium signalling and transcription *(Bähring, 2018)*. These proteins are also known to interact with Transient Receptor Potential (TRP) channels (*Dason* et al., *2012*) and are mainly expressed in the brain and antennae of *Aedes aegypti* (Matthews *et al*., 2016) and *Anopheles gambiae* (Pitts *et al*., 2011). These proteins are likely implicated in hearing, which directs our attention to the different mating behaviours between the two groups, swarming (mosquitoes), highly dependent on sound, and lekking (sandflies) where hearing is less relevant.

The four OGs showing increased diversification in both clades include *octopamine receptors* (OG0001113), *TBP-associated factor 3* (OG0001405), *qin* (OG0001557) and *WD-repeat protein outer segment 4* (OG0002006) gene families. In insects, octopamine is a key neuromodulator, often associated with odour-related behaviours (Claßen and Scholz, 2018), especially host-seeking (Tallon *et al*., 2020). Octopamine receptors are primarily expressed in the brain, ovaries (Matthews *et al*., 2016), and salivary glands (Ali, 1997). Amongst other functions, they are notable for modulating ovulation in mosquitoes (Lim *et al*., 2014) and vision in *Drosophila* (Suver *et al*., 2012). Furthermore, *Aedes aegypti* infected by dengue virus show dysregulation of octopaminergic signalling, causing reduced flight abilities and increased attraction to the odour of the host (Tallon *et al*., 2020). Based on these functions, we speculate that the diversification of octopamine receptors may have played an important role in the emergence of hematophagy, driving host-seeking behaviour in Diptera and, perhaps, in all biting insects. In *Drosophila*, WD-repeat protein outer segment 4 is involved in the regulation of the canonical Wingless pathway during development (Balmer et al. 2015), and in cilium assembly (Avidor-Reiss *et al*., 2004). In mosquitoes, it is expressed in antennal neurons (Matthews *et al*., 2016; Pitts *et al*., 2011) used for hearing and carbon dioxide perception (Montell and Zwiebel, 2016). Blood meals enable the transmission of pathogens from the host to the biting insect. Thus, we may expect a selective drive for the optimisation of the immune system of blood-feeding species. We have identified increased diversification within the *qin* family (OG0001557), which is involved in piwi-interacting (pi) RNA-mediated retrotransposon silencing by mRNA destabilisation (Zhang *et al*., 2011).This function can extend to viruses also (Varjak *et al*., 2018), which may explain the expansion of this family. The ingestion of blood requires enzymes for its digestion and detoxification. By structural similarity to other proteins, Qin has been proposed to have metal ion binding properties (Gramates *et al*., 2022), which may have a protective function.

In summary, our study has produced some intriguing insights on genetic changes that may have been permissive for the emergence of blood-feeding in mosquitoes and sandflies. Future work will be aimed at clarifying the function of the members of these gene families. Aditionally, we will address whether the evolutionary patterns we observed in these two clades can be generalised to other biting insects.

## Materials and methods

### Sampling and data pre-processing

Genomes, transcriptomes and proteomes for Culicomorpha (including mosquitoes and blackflies) and Psychodomorpha (including sandflies) were freely available on NCBI and were accessed in December 2021 (Benson *et al*., 2012). *De novo* assemblies of genomic and transcriptomic raw reads from the Sequence Read Archive (Leinonen *et al*., 2011) were assembled using MaSuRCA 4.0.5 (Zimin *et al*., 2013) and Trinity 2.13.2 (Haas *et al*., 2013). Proteomes were predicted using Augustus 3.2.3 (Keller *et al*., 2011) with either *Drosophila* or *Aedes* as a model species for genomic data, and TransDecoder 5.5.0 (https://github.com/TransDecoder/TransDecoder) for transcriptomic data. All proteomes were cleaned from duplicated sequences due to misassembly or sequencing errors by removing all sequences with more than 95% similarities using cd-hits 4.6.6 (Fu *et al*., 2012). Finally, proteomes were filtered for less than 30% missing Insecta BUSCO (v. 4.0.5) (Manni *et al*., 2021) genes the exception of a few species due to their ecological (Psychodidae: *Clogmia albipunctata*) or systematic relevance (Chironomidae: *Chironomus riparius*, Chaoboridae: *Chaoborus trivitattus*, Ceratopogonidae: *Culicoides sonorensis*). In total, 64 proteomes were analysed (see Supplementary Materials 1.0 S1).

### Phylogenetic inference

Species relationships were inferred using two different techniques: **(1)** Maximum Likelihood (ML) inference from a supermatrix and **(2)** a tree summarising method taking into account Incomplete Lineage Sorting (ILS) and gene flow which can interfere with phylogenetic inference methods (Degnan and Rosenberg, 2009).

Single-copy BUSCO genes were extracted from the previous completeness assessment and aligned using MAFFT 7.503 algorithm (Katoh and Standley, 2013). BUSCO assignments were corrected using a bespoke script, removing sequences divergent from more than twice the average standard deviation of the gene tree. Kept gene sequences were realigned as previously and trimmed using trimAl 1.2 (Capella-Gutierrez *et al*., 2009), allowing a maximum of half gaps in the final alignment. All gene-trimmed alignments were then concatenated with FASconCAT-G (https://github.com/PatrickKueck/FASconCAT-G) into a 641,089 amino acid positions across 64 species supermatrix. ML inference was carried out in version 2.1.4 of IQtree (Nguyen *et al*., 2015) under the best-fitting evolutionary model from ModelFinder (Kalyaanamoorthy *et al*., 2017). Ultra-Fast Bootstrap (UFB), as implemented in IQtree (Hoang *et al*., 2018), with 1000 replicates, was used to evaluate nodal support.

A second method, taking incomplete lineage sorting (ILS) into account was also used. Orthogroups were computed from the proteomes using OrthoFinder 2.0 (Emms and Kelly, 2019), identifying 63,621 gene trees for orthogroups with more than four sequences. The gene trees were then used to infer the species tree with the Accurate Species TRee ALgorithm for PaRalogs and Orthologs algorithm implemented in ASTRAL-Pro v.1.1.6 (Zhang *et al*., 2020). To encounter the lack of inference of tip branches length by the software, the topology of the tree was fixed in IQtree using the previous single-copy BUSCO genes supermatrix. Node supports of the tree are shown as Local Posterior Probabilities (LocalPP) as described in (Zhang *et al*., 2020).

### Estimation of divergence times

Divergence times among the two inferred trees were inferred with the penalised likelihood method implemented in r8s v. 1.81 (Sanderson, 2003), using a 260 Mya split between *Drosophila melanogaster* and *Anopheles gambiae* as a calibration point (Wiegmann *et al*., 2011).

### CAFE analysis

We then used a birth/death model with a gamma distribution implemented in CAFE 5 (Mendes *et al*., 2021) to detect changes in the rate of evolution of individual gene families along the trees. The parameters of the birth-death model were estimated using a subset of OGs with less than 200 genes. Because of software limitations for this analysis, we removed the 25 largest gene families with more than a thousand genes. All orthogroups have been annotated under eggNOG-mapper version 2 (Cantalapiedra *et al*., 2021)..

### Ancestral state reconstruction of blood-feeding behaviour

All subsequent analyses were carried out in RStudio using R version 4.3 (R Core Team, 2024). Hematophagy was encoded as a discrete trait (0 for non-biting and 1 for biting). The ancestral state has been inferred among both previous ultrametric trees using the phylogenetic ridge regression implemented in the R package *RRphylo* (Castiglione *et al*., 2020). Ancestral states were then plotted on the trees with *ggtree* (Yu *et al*., 2017). Nodes A and B (Figure 1) were used for the emergence of hematophagy in both clades for the following analysis.

### Selection of gene families of interest and Gene tree-species tree reconciliations

Orthogroups with a significant (p-value<0.05) change in their rate of evolution for both nodes A and B in both trees were filtered out. Gene tree-species tree reconciliation of the five gene families showing changes in their rate of evolution at both nodes A and B was performed in GeneRax v. 1.1.0 (Morel *et al*., 2020) using duplication and loss models. Reconciled trees were visualised using ThirdKind 3.2.2 (Penel *et al*., 2022). The effect of hematophagous behaviour (blood-feeding *vs*. non-blood feeding) on the number of genes in these gene families has been tested using an ANOVA corrected for the phylogeny and implemented in the R package *geiger* version 2.0.11 (Pennell *et al*., 2014). To better describe these gene families, *Drosophila melanogaster* sequences for the five gene families of interest were aligned against its reference proteome in FlyBase release FB2024_01 (Gramates *et al*., 2022) using a Basic Local Alignment Search Tool (BLAST). BLAST of *Anopheles gambiae* and *Aedes aegypti* sequences was also performed in VectorBase release 67 (Giraldo-Calderón *et al*., 2015).

## Acknowledgements

This work is supported by a University Research Fellowship (UF160226 and URF/R/221011) to R Feuda. JD is supported by a PhD Scholarship from the University of Leicester. RF and JD also acknowledge Dr Mattia Giacomelli for providing the script for BUSCO orthology correction. This research used the ALICE High-Performance Computing Facility at the University of Leicester.

## Author contributions

JD and RF conceived the study. JD conducted the experiment. JD analysed the data. JD and RF wrote the draft manuscript. All authors contributed to and reviewed the final manuscript.

## Data Accessibility

All code is available at: https://github.com/juliendevi/phylogenomics_mosquitoes.

## Supplementary 2.0: additional figures

**Figure S1. Phylogenetic inference using ASTRAL and maximum likelihood methods**. Incongruence between methods on both trees are shown in red. Mosquitoes and sand-flies (Psychodidae) are identified in orange. A - Maximum likelihood inference using a supermatrix of single-copy BUSCO genes, ultrafast boostraps diffrent from 100 are shown on the nodes. B - ASTRAL inference using 63,621 gene trees, Local Posterior Probabilities different from 1 are shown on the nodes.

**Figure S2. Phylogenetic and ancestral trait inference of blood feeding behaviour in Culicomorpha and Psychidomorpha**. Ultrametric BUSCO tree obtained from a supermatrix with ML inference using IQ-tree (Minh et al., 2020) and divergence times computed in r8s (Sanderson, 2003). Colour of the nodes shows the probabilities for ancestral behaviour (blood feeding [red] or non-blood feeding [blue]) from a phylogenetic ridge regression obtained with RRphylo (Castiglione et al., 2020). Nodes A and B respectively represent our points of interest with emergence of hematophagy in mosquitoes and sand-flies.

**Figure S3. ASTRAL gene tree-species tree reconciliation of gene families showing significant changes in gene number in the LCMA and LCSA**. On the ASTRAL topology are shown for each nodes the number of genes inferred (CAFE/GeneRax), duplications (in green) and losses (in red) events inferred by CAFE. Significant events are highlighted by a star.

**Figure S4. BUSCO gene tree-species tree reconciliation of gene families showing significant changes in gene number in the LCMA and LCSA**. On the BUSCO topology are shown for each nodes the number of genes inferred (CAFE/GeneRax), duplications (in green) and losses (in red) events inferred by CAFE. Significant events are highlighted by a star.

