## Supplementary figures and images for "Hematophagy generates a convergent genomic signature in mosquitoes and sandflies"

### FigureS1

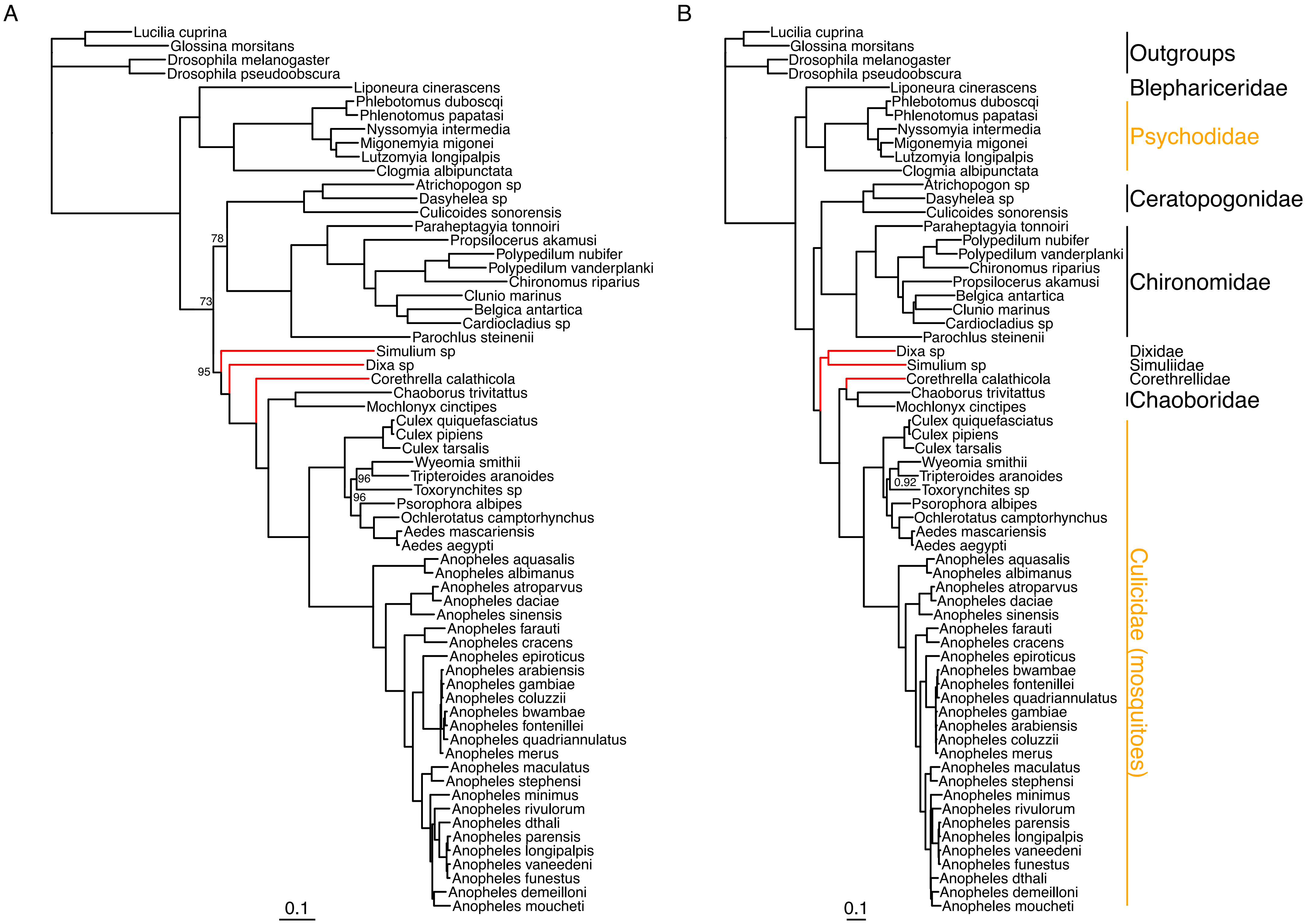

### FigureS2

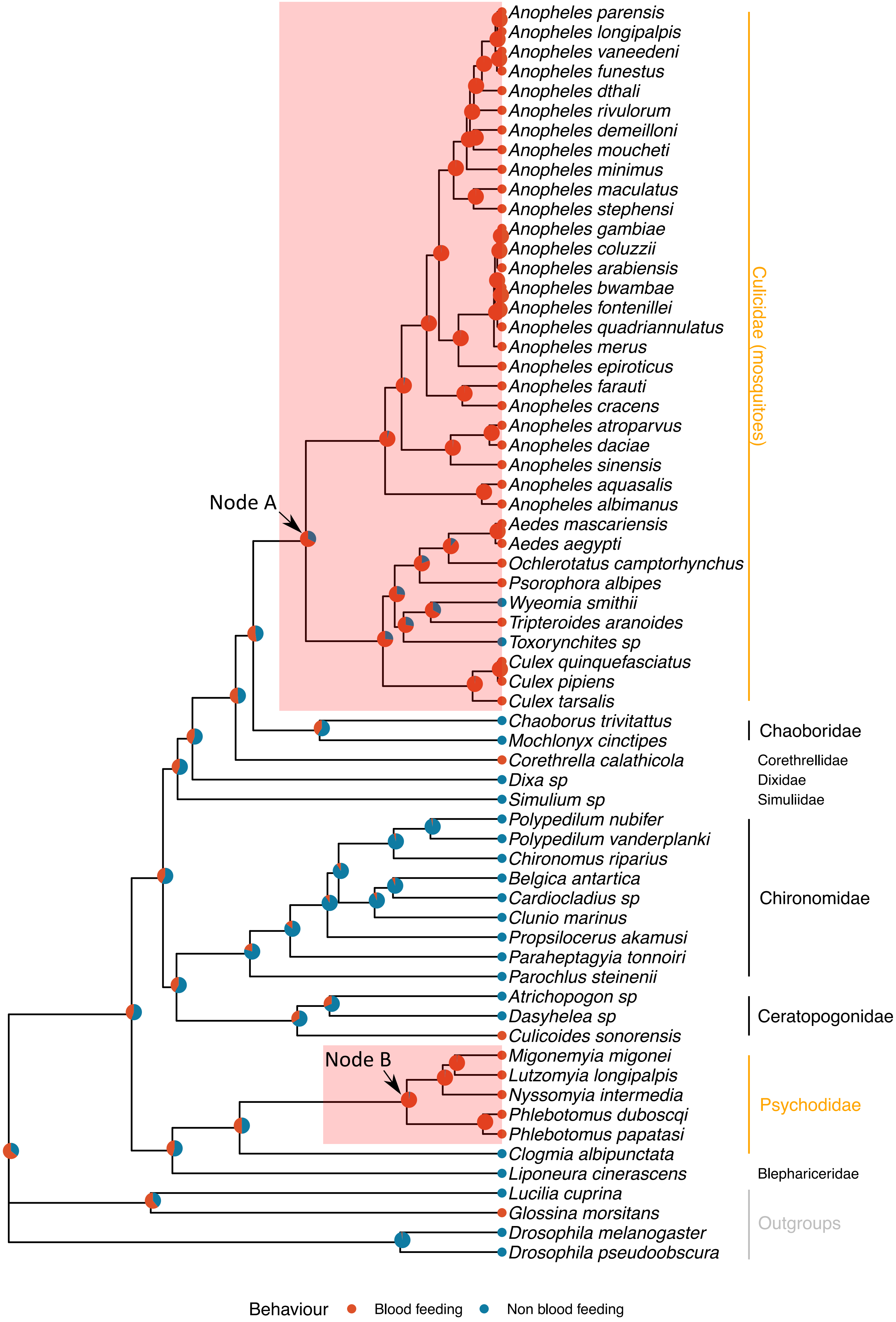

### FigureS3

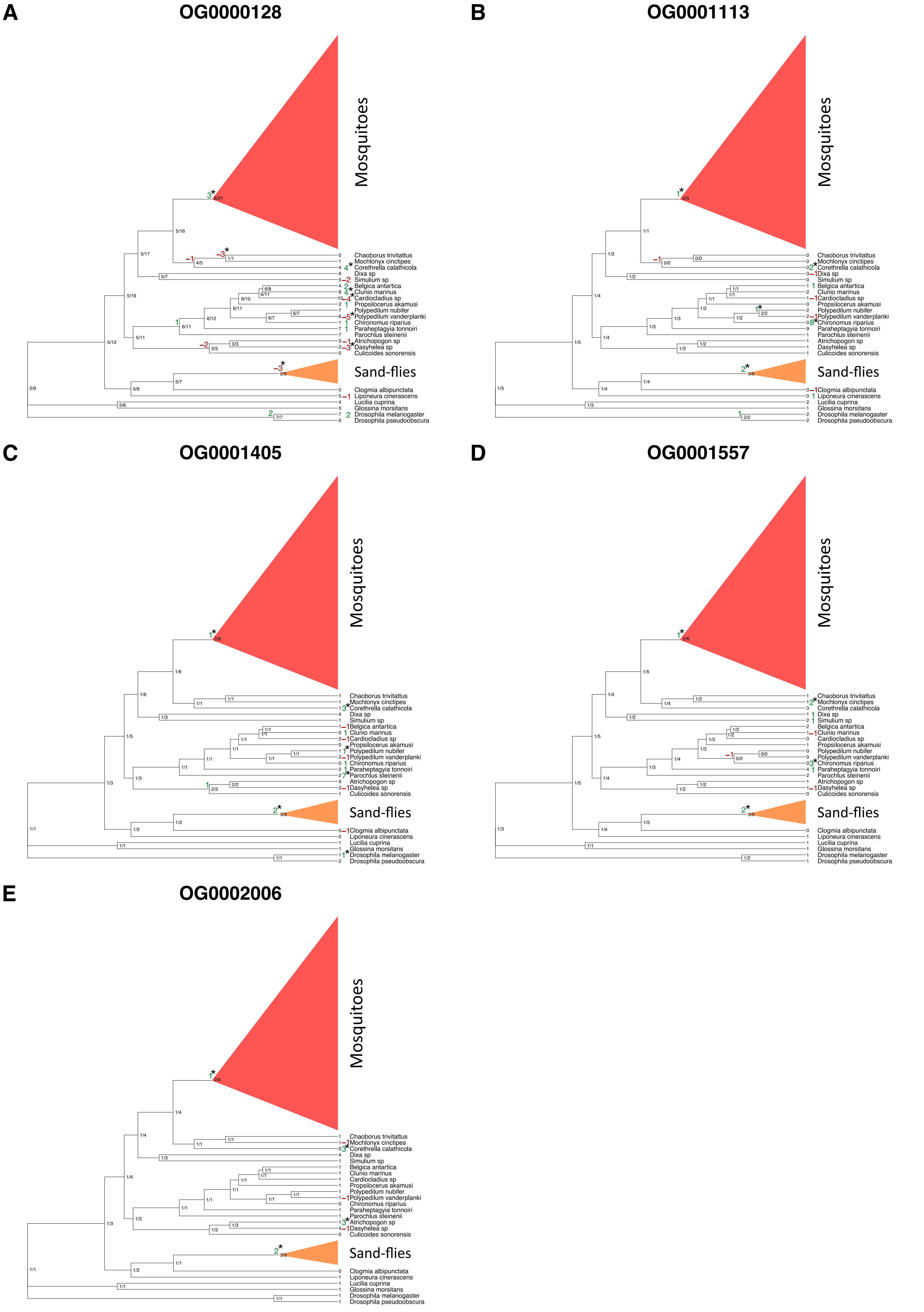

### FigureS4

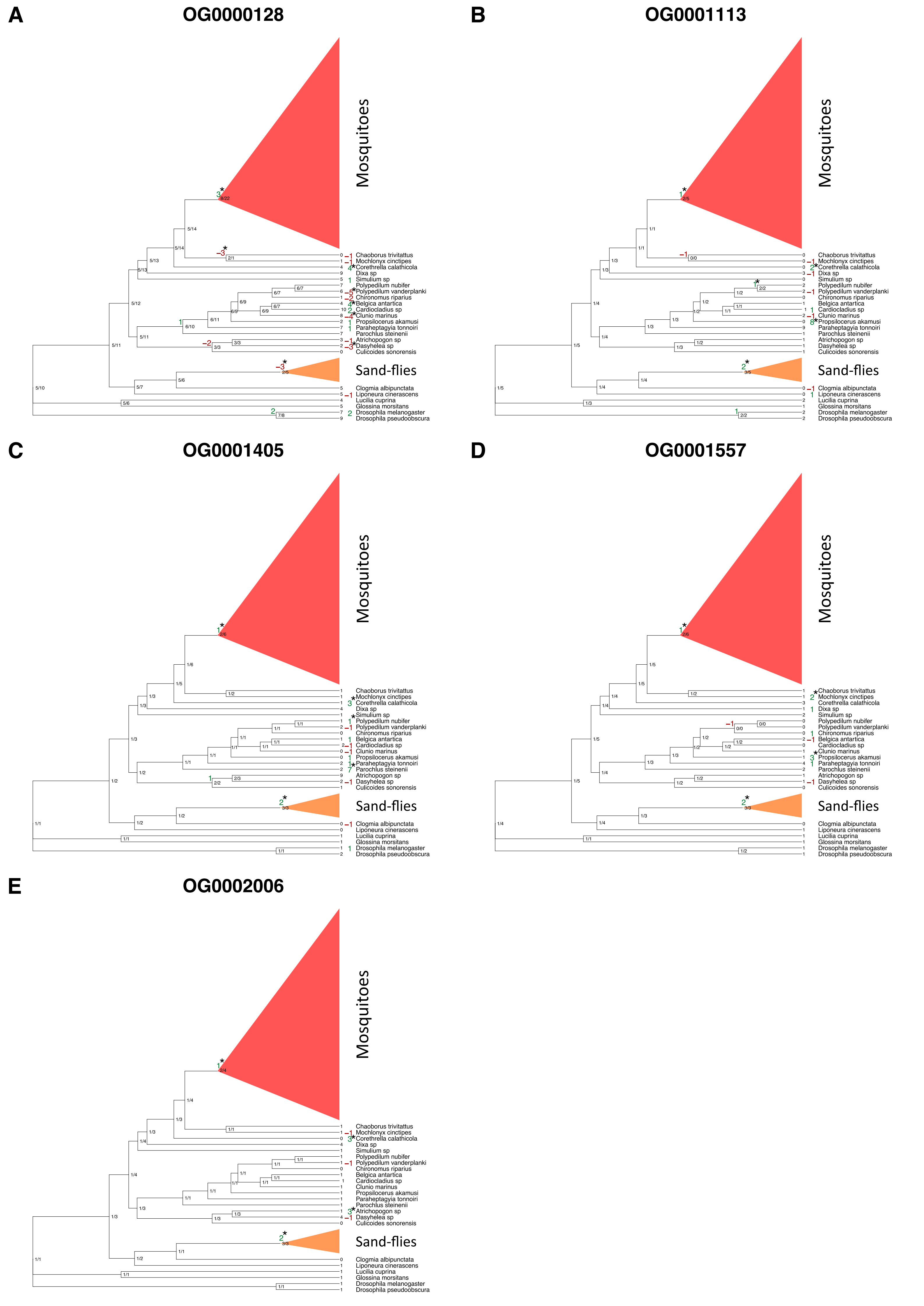
